# Elucidation of Genome-wide Understudied Proteins targeted by PROTAC-induced degradation using Interpretable Machine Learning

**DOI:** 10.1101/2023.02.23.529828

**Authors:** Li Xie, Lei Xie

**Affiliations:** Department of Computer Science, Hunter College, The City University of New York, New York, 10065, USA; Ph.D. Program in Computer Science, The Graduate Center, The City University of New York, New York, 10016, USA; Helen and Robert Appel Alzheimer’s Disease Research Institute, Feil Family Brain & Mind Research Institute, Weill Cornell Medicine, Cornell University, New York, 10021, USA

## Abstract

Proteolysis-targeting chimeras (PROTACs) are hetero-bifunctional molecules. They induce the degradation of a target protein by recruiting an E3 ligase to the target. The PROTAC can inactivate disease-related genes that are considered as understudied, thus has a great potential to be a new type of therapy for the treatment of incurable diseases. However, only hundreds of proteins have been experimentally tested if they are amenable to the PROTACs. It remains elusive what other proteins can be targeted by the PROTAC in the entire human genome. For the first time, we have developed an interpretable machine learning model PrePROTAC, which is based on a transformer-based protein sequence descriptor and random forest classification to predict genome-wide PROTAC-induced targets degradable by CRBN, one of the E3 ligases. In the benchmark studies, PrePROTAC achieved ROC-AUC of 0.81, PR-AUC of 0.84, and over 40% sensitivity at a false positive rate of 0.05, respectively. Furthermore, we developed an embedding SHapley Additive exPlanations (eSHAP) method to identify positions in the protein structure, which play key roles in the PROTAC activity. The key residues identified were consistent with our existing knowledge. We applied PrePROTAC to identify more than 600 novel understudied proteins that are potentially degradable by CRBN, and proposed PROTAC compounds for three novel drug targets associated with Alzheimer’s disease.

**Author Summary:** Many human diseases remain incurable because disease-causing genes cannot by selectively and effectively targeted by small molecules. Proteolysis-targeting chimera (PROTAC), an organic compound that binds to both a target and a degradation-mediating E3 ligase, has emerged as a promising approach to selectively target disease-driving genes that are not druggable by small molecules. Nevertheless, not all of proteins can be accommodated by E3 ligases, and be effectively degraded. Knowledge on the degradability of a protein will be crucial for the design of PROTACs. However, only hundreds of proteins have been experimentally tested if they are amenable to the PROTACs. It remains elusive what other proteins can be targeted by the PROTAC in the entire human genome. In this paper, we propose an intepretable machine learning model PrePROTAC that takes advantage of powerful protein language modeling. PrePROTAC achieves high accuracy when evaluated by an external dataset which comes from different gene families from the proteins in the training data, suggesting the generalizability of PrePROTAC. We apply PrePROTAC to the human genome, and identify more than 600 understudied proteins that are potentially responsive to the PROTAC. Furthermore, we design three PROTAC compounds for novel drug targets associated with Alzheimer’s disease.

## 1 Introduction

Despite rapid progress in the development of small-molecule drugs, traditional drug discovery is limited by the requirement of efficient inhibitions on functional binding sites in protein targets. However, many of disease-associated proteins neither have suitable binding pockets for small molecule drugs nor bind inhibitors with considerable affinity [1]. It is estimated that only 2-5% of the human genome are druggable by small molecule drugs [1, 2]. For example, in the anti-cancer drug discovery, major cancer drug targets belong to serine/threonine/tyrosine kinases, growth factor receptors and GPCRs. Other classes of cancer-relevant proteins, like phosphatases, transcription factors, and RAS family members, have difficulties to be inhibited by small molecules and are considered to be understudied. Thus, it is necessary to seek alternative approaches to target these proteins. Several types of biological drugs, including peptides, antibodies, modified nucleic acids, and vaccines are developed to target these proteins. However, the larger size of the biological drugs limits their delivery mode and makes it more challenging to alter intracellular targets than small molecule drugs. [3].

Different from small molecule and biological drugs, protein-targeting chimeric (PROTAC) molecules are hetero-bifunctional molecules that have a small molecule binding moiety for a target protein (i.e., warhead) and an E3 ubiquitin ligase-recruiting moiety coupling by a chemical linker. With the two binding moieties, PROTACs recruit an E3 ubiquitin ligase to a targeted protein and induce the degradation of the whole targeted proteins in the ubiquitin–proteasome system [4, 5, 6]. Thus, even when the binding site of a PROTAC is not on the functional domain for some targeted proteins like understudied proteins, PROTACs can still shut down their functions by the degradation of the whole protein [7, 8, 9, 10]. This feature gives PROTACs potentials to overcome the limitation of small molecule drugs when targeting understudied proteins, such as transcription factors, scaffolding and regulatory proteins. [11, 12, 13].

PROTACs can increase both selectivity and efficacy of targeted therapies. In the PROTACs-induced degradation, a targeted protein, a PROTAC and an E3 ligase form a ternary complex structure. The selectivity of PROTACs is not only determined by the warhead, but also by the protein-protein interaction between the E3 ligase and the targeted protein as well as the stability of the ternary complex structure [5, 12, 13]. Even with a promiscuous warhead, PROTACs can still display enhanced target selectivity compared with small molecule inhibitors [14]. Moreover, low-affinity ligands or ligands failing to modulate targeted functions are usually considered as ineffectual ligands in the traditional drug discovery but they still can be used to design PROTACs to induce the degradation for target proteins [15]. Meanwhile, for multi-functional proteins, small molecule or biological drugs can only inhibit the occupied functional sites, but PROTACs can target all functions by eliminating the whole proteins. Thus PROTACs can increase pharmaceutical efficiencies. For example, a PROTAC designed for receptor tyrosine kinases was shown to inhibit both cell proliferation and downstream signaling [16], and focal adhesion kinase PROTAC affected both tumor invasion and migration [17, 18]. Recently, PROTAC-induced degradation has been applied to multi-protein complexes to modulate multiple functions on the whole complex upon the degradation on one component, leading to destabilization and subsequent degradation of other complex components [19]. Such ability would greatly increase the current drug target space or facilitate addressing drug resistance problems by targeting the multi-component complex system. There are several successful cases in the degradation of epigenetic proteins, including epigenetic reader protein BRD4 [20, 21, 22], epigenetic eraser proteins [23] and others [15, 24, 25], resulting in inhibitions on both enzymatic and scaffolding functions for epigenetic protein complexes. Studies on polycomb repressive complex 2 (PRC2) [26, 27, 28] also showed that PROTAC targeting of EED can cause EED, EZH2, and SUZ12 protein loss, leading to the perturbation of histone methylation and eliminated cell proliferation.

Another advantage of PROTACs induced degradation over traditional inhibition is the ability to address acquired drug resistances [16, 29, 30, 31]. For example, BKT inhibitor ibrutinib-derived PROTAC can induce degradation of both wild-type and C481S-mutant BTK to overcome ibrutinib resistance [30]. This study suggested that PROTACs can circumvent resistance mechanisms affecting parent inhibitors. Several studies found that PROTACs could overcome the drug resistance to original targets by targeting other proteins. For example, an RTK inhibitor lapatinib conjugated with VHL ligand produced a PROTAC that is capable to induce the degradation of membrane-bound wild-type EGFR as well as disease-relevant EGFR mutants [16].

These advantages intrinsic to the mechanism of action of PROTACs make the PROTAC-induced degradation a very promising therapeutic modality with the potential to reduce drug exposure requirements, enhance drug selectivity, circumvent drug resistance mechanism, and target the understudied disease genes. More and more researches focus on this field, including both experimental and computational approaches. However, computational-based design for PROTACs is still limited to a small number of published ternary complex structures, and mainly focuses on the modeling of ternary complex structures with a pre-designed PROTAC through PRosettaC [32], Rosetta [33], RosettaDock [34], docking combined with molecular dynamic simulations [35, 36, 37, 38] and molecular modeling on linkers [5, 39]. No studies have been conducted to explore understudied proteins that are amenable to PROTACs in the entire human genome. In this work, we have developed an interpretable machine learning model PrePROTAC to predict degradable proteins targeted by PROTACs using embedding features extracting from protein sequences. PrePROTAC is the first model of the kind, and it allows us to elucidate understudied proteins whose degradability can be induced by PROTACs on a genome scale. We have identified about 615 new proteins that are responsible for multiple diseases can be targeted by the PROTACs. Furthermore, we screened the PROTACs for three novel drug targets for Alzheimer’s disease.

## 2 Results and discussion

### 2.1 Overview of PrePROTAC

The whole process of our PrePROTAC model was shown in Fig 1a. Starting from a protein sequence, a sequence embedding was generated using protein language model ESM. The sequence embedding was used as features to train a Random Forest (RF) classification model to predict the degradability of this protein targeted by CRBN. Meanwhile, an eSHAP interpretative model was developed to identify the key residues on the target protein by calculating the difference of the selected features between the original sequence and its single-residue mutated sequences. The details of the model training, validation and eSHAP analysis can be found in the method section. The trained PrePROTAC model was applied to the whole human proteome. A threshold of 0.9 was used to select the proteins which have the potential to be degraded by CRBN when induced by PROTACs. As shown in Fig 1b, Autodock Vina, Zdock and Rosseta docking were used to screen the PROTACs and build ternary complex structures for selected proteins. eSHAP analysis was used to identify the residues which might play important roles in the degradation.

**Fig 1.**
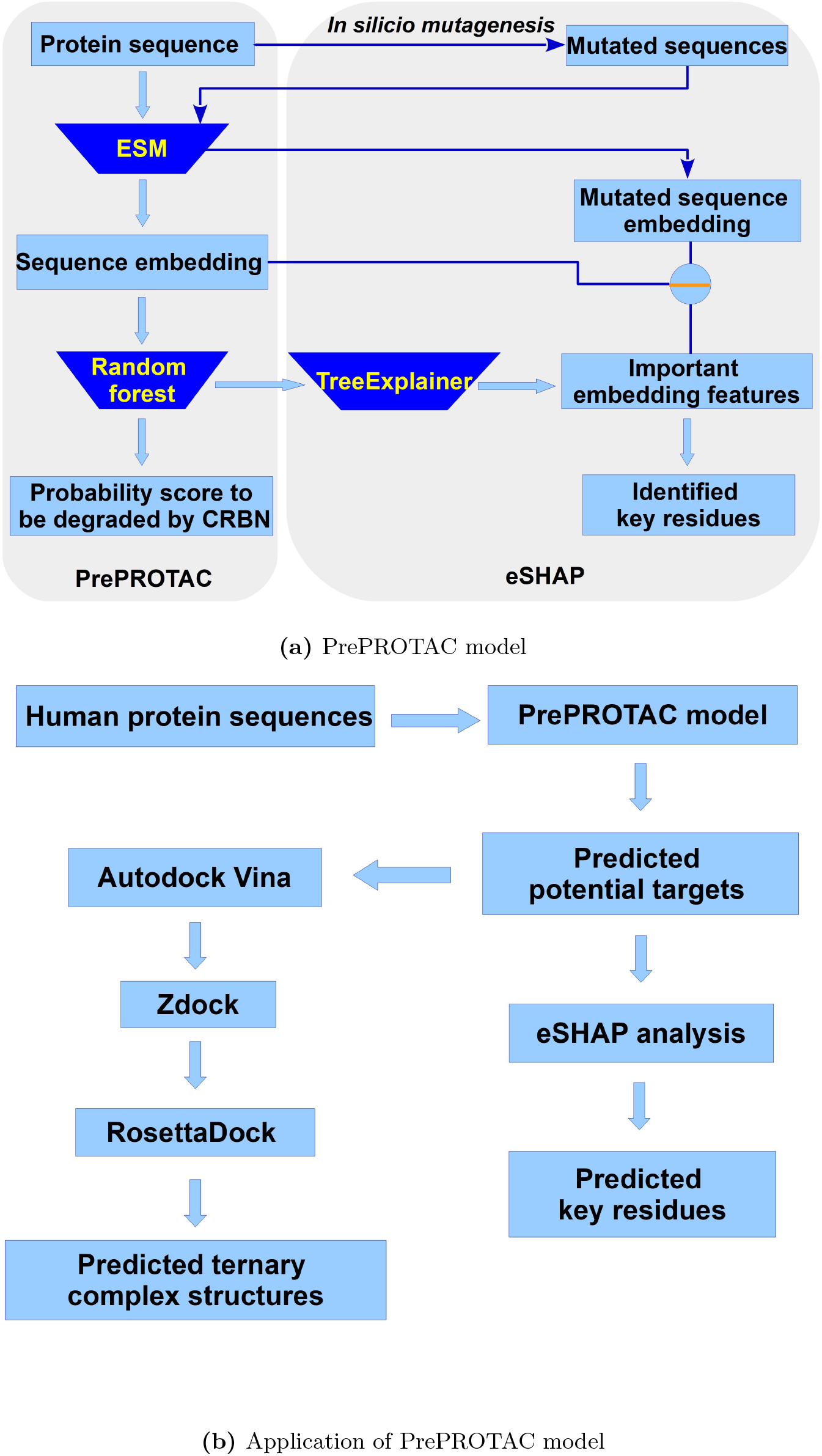
Overview of the PrePROTAC mdoel and application process.

### 2.2 Comparison between Random Forest and Gradient Boost Tree classification methods with Ifeatures, D-script contact features and ESM features

Because only less than 500 proteins had annotated degradability information, but protein sequences were usually represeted by a high-dimensional vector, we selected Random Forest (RF) and Gradient Boost Tree (GBT) to train the prediction model because they are less prone to over-fitting for high-dimensional data with a small number of samples. In order to obtain optimal performances, hyper-parameters used in the RF and GBT models with different features were tuned through 5-fold grid search cross validations and the optimized hyper-parameters for different models and features were listed in S1 Table and S2 Table.

With the tuned hyper-parameters, RF and GBT classification models corresponding to 21 Ifeature descriptors[40], D-Script contact feature[41] and ESM feature[42] were trained to fit the protein kinase degradation data. Repeated Stratified 5-Fold cross validation method was applied to validate these models. Here the Stratified 5-fold cross validation was repeated twice to get a more accurate estimate of the performance distribution of the model. Average ROC-AUC scores for total 42 classification models were shown in S1 Fig.

For the 21 Ifeature descriptors, the one with the best ROC-AUC score in each group was selected and compared with D-script and ESM features. Models based on TPC, GTPC, Geary, CTDD, CTriad and QSOrder had the best performances compared with the other descriptors in the same group. As shown in Fig 2, the RF model with ESM feature had the best performance when evaluated by average ROC-AUC scores and average precision scores. Performance of the GBT model with ESM feature was comparable with the RF model. Their ROC-AUCs were 0.764 and 0.765, and precision scores were 0.74 and 0.75, respectively. Among TPC, GTPC, Geary, CTDD, CTriad, QSOrder, D-script contact feature and ESM feature, ESM feature outperformed all others. In the follow-up studies, the ESM feature was exclusively used. ROC curves, false positive rate curves, and precision-recall curves for the RF and GBT with TPC, GTPC, Geary, CTDD, CTriad, QSOrder, D-script contact feature and ESM feature were shown in S2 Fig-S7 Fig.

**Fig 2.**
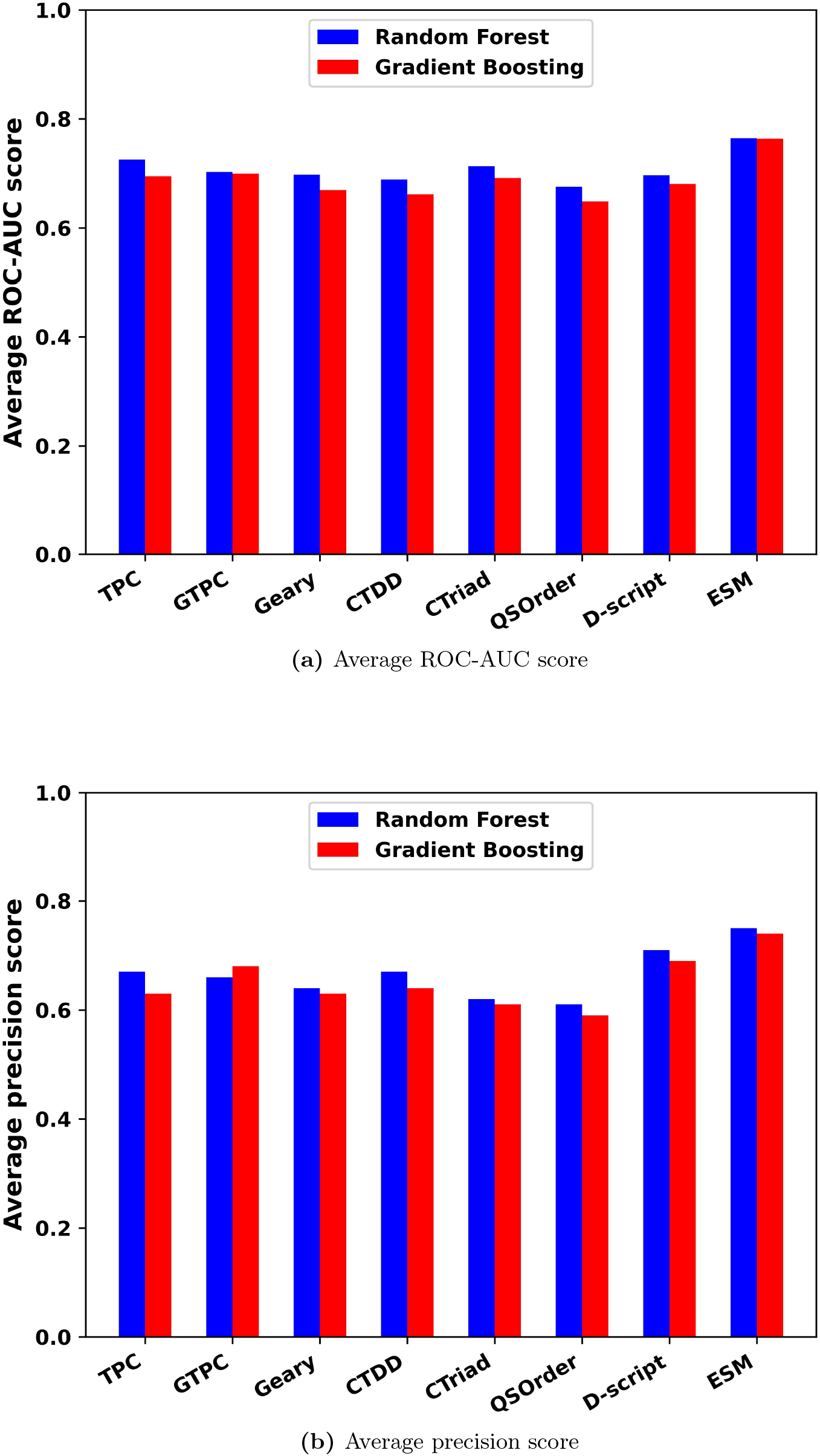
Average ROC-AUC scores and precision scores for RF and GBT classification models with 8 features. (a) Average ROC-AUC score. (b) Average precision score.

We further evaluate the performance of models using an external test data set in which no proteins are protein kinases (See Method for details). When using the randomly selected proteins in the same SCOP fold but different SCOP super-families from the degradable proteins as the negative samples, the ROC-AUC score for the prediction is about 0.71, as shown in Fig 3. However, the performance of the GBT model was significantly worse than the RF model. The average ROC-AUC is 0.64. The average precision score for the GBT model was also worse than that of the RF model, as 0.61 compared with 0.68.

**Fig 3.**
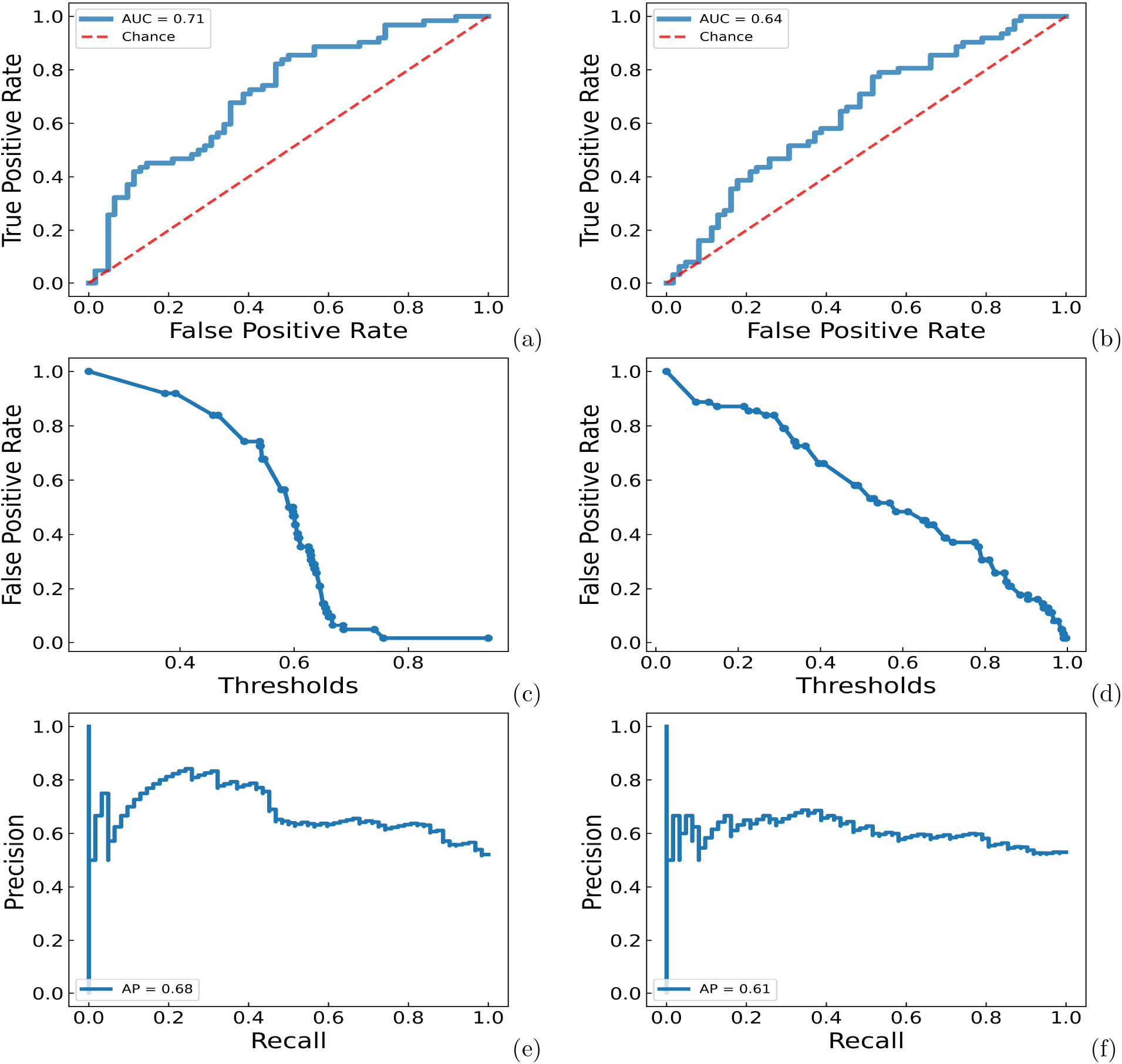
ROC-AUC, false positive rate-threshold, precision-recall curves for RF and GBT models on the test set with SCOP proteins as negative samples. (a) ROC-AUC curve for the RF model. (b) ROC-AUC curve for the GBT model. (c) false positive rate-threshold curve for the RF model. (d) false positive rate-threshold curve for the GBT model. (e) precision-recall curve for the RF model. (f) precision-recall curve for the GBT model.

In order to investigate whether the performance could be improved when adding other features with ESM feature, TPC, GTPC, Geary, CTDD, CTriad, QSOrder and Dscript features were combined with ESM feature. Hyper-parameters for the RF models with these combined features were trained and 5-fold cross validations of these models were performed on the training set. Average ROC-AUC scores and average precision scores were shown in S8 Fig. Even compared with these combined features, performances of the ESM feature alone was still the best. Simply adding these features to the ESM feature could not improve the performance.

### 2.3 Performance of RF and GBT ensemble models

In order to investigate whether the performance could be improved when combining the RF and GBT models, two different ensemble methods were used: one is the soft voting method and the other one is the consensus method. In the soft voting method, average probability scores of the RF and GBT models were used to predict the degradation of proteins. For the consensus method, only when both models agree positive, the result will be positive. Otherwise, the result will be negative. Five fold cross validation method was used to evaluated the performances of the soft voting model and consensus model on the training set. Average ROC-AUC, false positive rate-threshold, precision-recall curves for them were shown in S9 Fig. Both of the ensemble models had similar performances with the RF model. When the ensemble models were evaluated using the external test set, their performances were better than the GBT model, but still worse than the RF model, as shown in S10 Fig. In general, for all these tested models based on two different classification methods with 23 different features, the RF classification model with ESM feature got the best performance and would be used in the subsequent studies.

### 2.4 eSHAP analysis identified key residues for the PROTAC activity of protein kinases

eSHAP analysis was applied to protein kinases in the training set and identified the key positions for the PROTAC activity. In order to further understand the functional roles of these key positions, these key positions across the protein kinase family were mapped to the structures of these protein kinases. Structure-based multiple sequence alignments (MSAs) were obtained from 497 human protein kinase domains [43]. Identified key positions of the protein kinases were mapped on the MSA. A key-position only MSA was formed by chopping other residues. Then, these protein kinases were classified into 12 subgroups according to the key-position only MSA. For a better visualization of such classification, a key-position MSA related protein kinase tree was created using the maximum likelihood algorithm in the software MegaX [44] and visualized with the webserver iTOL [45]. As shown in Fig 4, different from typical protein kinase phylogenetic trees built from the MSA of whole protein kinase domains [43][46][47], branches in this tree generated from the key-position only MSA contained protein kinases from different protein kinase subfamilies.

**Fig 4.**
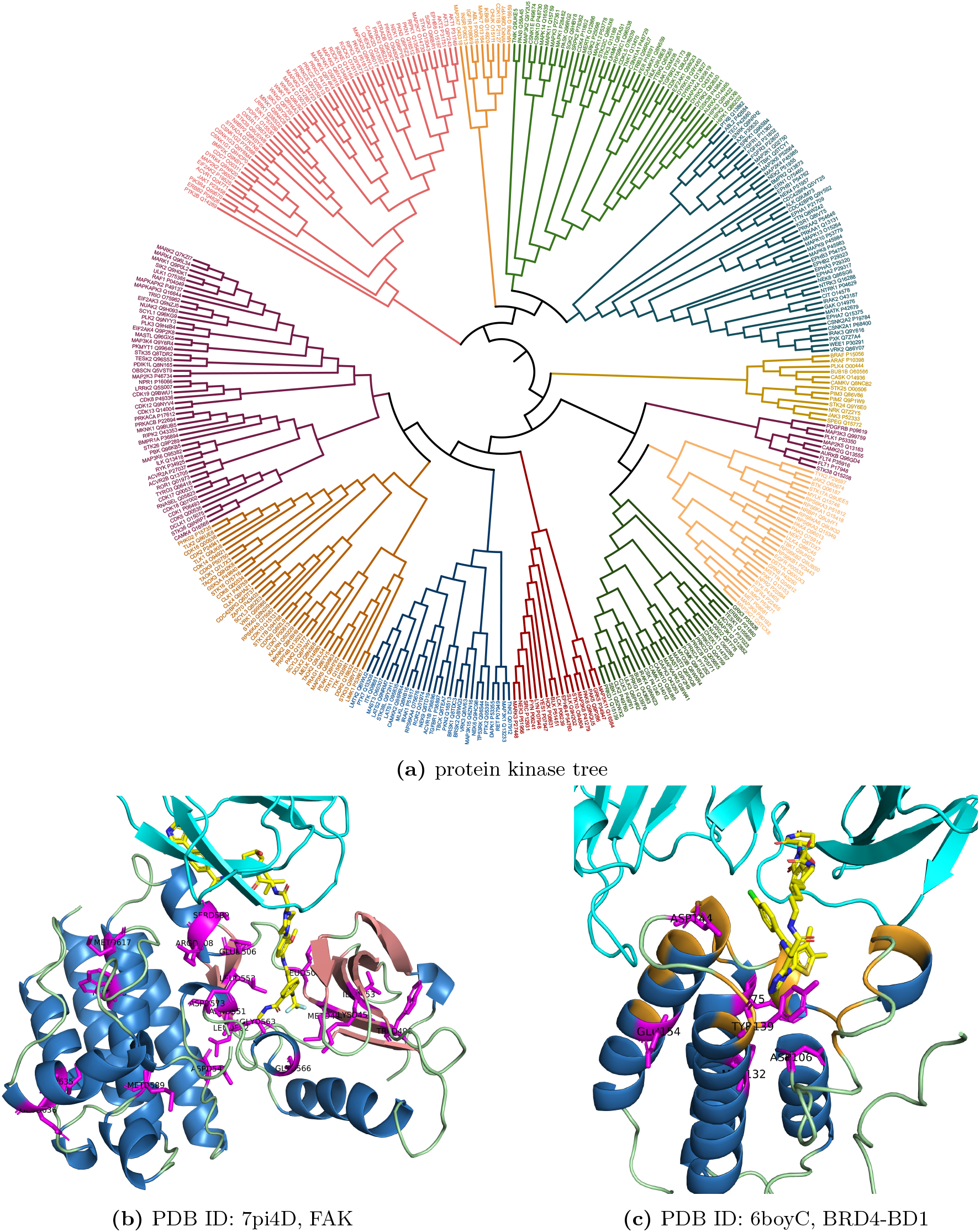
Protein kinase tree based on the key-position only alignments and structural mapping of the key positions for FAK and BRD4. Pink represents strands, blue represents helices, green represents loop regions, yellow represents co-crystalized ligand and magenta represents the top ranked positions, orange represents binding sites.

In these protein kinases, FAK has a ternary complex structure with CRBN and PROTAC. The key positions identified for FAK were mapped on the complex structure (PDB ID: 7pi4). As shown in Fig 4, most of these key positions were found in the PROTAC binding pocket or the interface between FAK and CRBN, including Ile453 and Lys454 on the *β*-strand 3, Trp496, Met499 and Leu501 on the strand 5, Glu506, Arg508 and Ser509 on the D helix, Asp546 and Asn551 on the catalytic loop, Leu553, Leu562 and Gly563 on the N terminal of the activation loop (ALN), Gly566 on the DFG motif that is the part of ALN, Asp573 and Met589 on the C terminal of the activation loop (ALC). Trp613 and Met617 on the F helix, Ile635 and Glu636 on the G helix. All of these segments are of functional importance for the protein kinase. Lys454 on the *β*-strand 3 is the catalytic residue which usually interacts with the *α-* and *β*-phosphates of ATP. Met499 is the gatekeeper residue which influence the accessibility to a buried region at the end of the ATP binding pocket and controls inhibitor sensitivity of protein kinases. Other residues on the *β*-strand 3 and 5 forms part of the buried region in the ATP binding pocket. Asp 546 and Asn551 are located on the catalytic loop and close to the gamma-phosphate group of ATP, so they are directly related to catalytic functions of FAK. Leu553, Leu562 and Gly563 locate on the ALN. Gly566 is one of the DFG residues. ALC contains the APE motif which can stabilize the activation loop by docking to the F helix. Met589 is next to the APE motif. GLY566 and Asp573 on the activation loop could bind the substrates and thus promote the catalysis. F helix is a highly hydrophobic segment and serves as a stable anchor for the most other motifs in the C-lobe including the catalytic loop and the activation Loop through hydrophobic contacts with them. G helix is a solvent exposed helix and a part of GHI-domain. This domain can interact with many substrate proteins and regulatory proteins and act as allosteric sites. Key positions identified for other protein kinases also locate on these functional important segments. Representatives of the positive and negative samples for each protein kinase subgroups were selected according to the key-position only MSA. The structural mapping of their key positions and the key-position only MSA were shown in S11 Fig-S22 Fig.

Except for FAK, the BD1 domain of BRD4 (PDB ID: 6boy) also form a ternary complex structures with CRBN and its PROTAC. The key positions for this protein were also identified and mapped on their complex structures to show the functional roles for these key positions. BRD4 is one of the members in BET family, which regulate the expression of many immunity-associated genes and pathways. It is also a well-studied target protein for PROTAC-induced degradation [34] [20][48]. The BD1 domain of BRD4 can be degraded by CRBN through PROTAC-induced degradation. The top five key positions on BD1 were shown in the ternary complex structures with CRBN and PROTAC. Cys136, Tyr139 and Asp144 are the ligand binding residues. Other top positions are also close to the ligand binding pockets.

In summary, the key residues identified by eSHAP analysis were supported by exisiting biological knowledge. They provided additional validations to PrePROTAC.

### 2.5 Application on human disease-associated understudied proteome

In order to apply PrePROTAC to unseen proteins on a genome scale, we further developed a soft voting classification model based on the ten models with equal weight (See Methods for details). When the model was evaluated by the testing data from all samples, ROC-AUC curves, average false positive rate-threshold curve and precision-recall curve for the ten models were shown in Fig 5. Based on the correlation between average false positive rate and threshold from ten validation sets, if the threshold is set as 0.85, the false positive rate would be 0.004, which means only one of the negative sample was predicted as positive since there were 241 negative samples in the data set. In order to select the potential degradable proteins with high confidence, the threshold of 0.90 would be used to select positive hits in the prediction. We applied PrePROTAC to 20,504 human proteins (see Method for selection procedure). The probability of these proteins to be degraded by CRBN were predicted by the soft voting model and the distribution of probability scores were presented in the S23 Fig.

**Fig 5.**
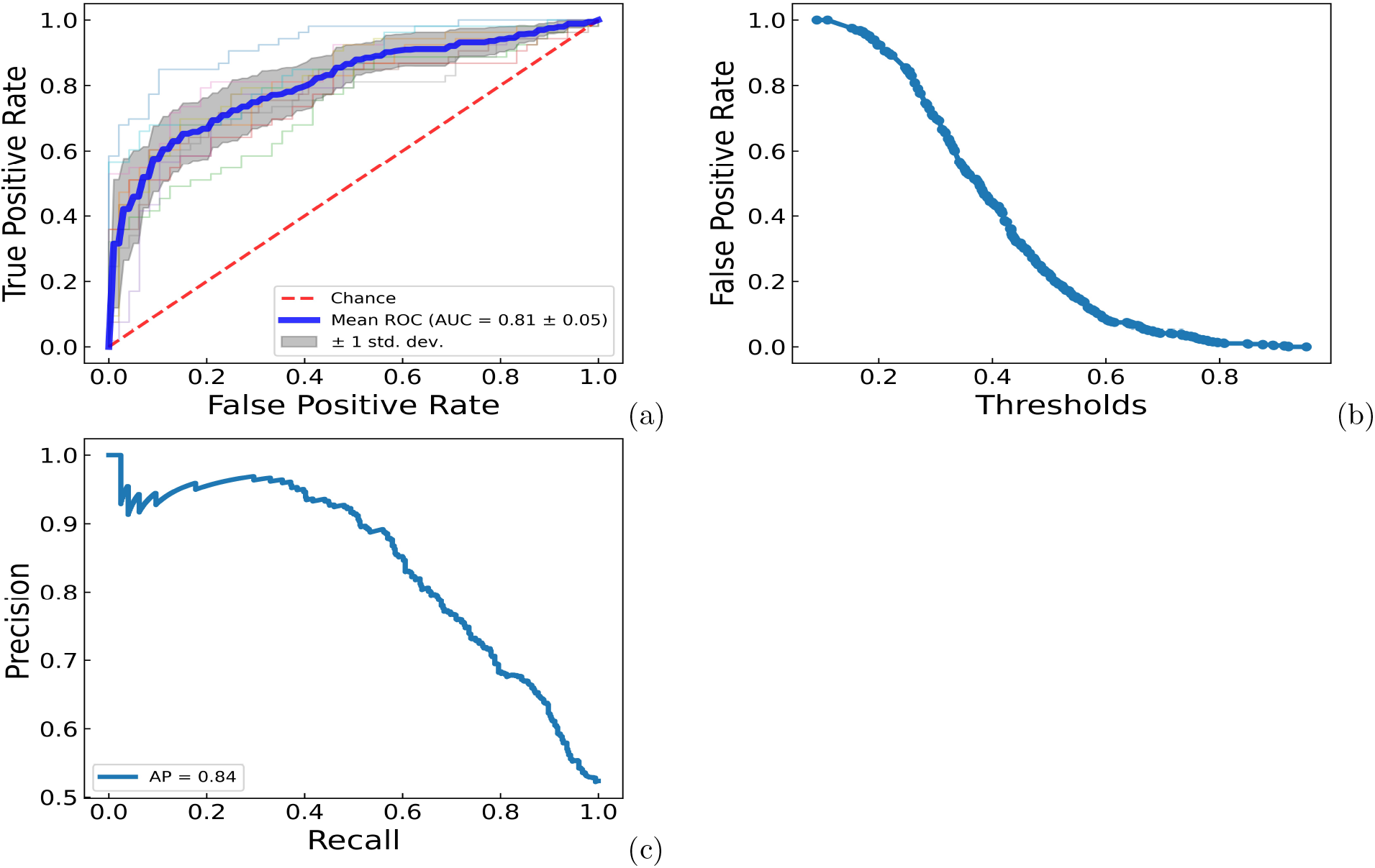
ROC-AUC curve, average false positive rate-threshold curve and average precision-recall curve for ten ESM RF models. a) ROC-AUC curve. b) Average false positive rate-threahold curve. c) Average precision-recall curve.

It is well known that only a subset of the human genome that is considered druggable with studies on drug-like small molecules. There is only limited information for the understudied disease proteins [49]. Hence, we built an understudied human protein database by removing the druggable proteins (Tclinic and Tchem) in Pharos [50] and Casas’s druggable proteins [51] from human disease associated genome database [52] and applied our method to predict the probability of these understudied disease proteins to be degraded by CRBN in the PROTAC-induced degradation. The understudied disease protein data set include 12,475 human proteins and these proteins were ranked according to their probability scores. 615 of them were predicted to be degradable by CRBN with the help of PROTAC binding. Information about these proteins and their predicted probability scores were listed in the supplementary table S3.

In order to help design PROTACs for the proteins predicted positive by our PrePROTAC model, protein-protein docking and protein-ligand docking were used to look for PROTACs which could bind to both CRBN and selected target proteins. Three Alzheimer’s diseases related proteins were selected according to their PrePROTAC prediction scores, including Ski-like protein fragment (UniProt ID: P12757), death domain-associated protein 6 (UniProt ID: Q9UER7) and activating signal cointegrator 1 complex subunit 2 (UniProt ID: Q9H1I8). These proteins are highly expressed in the brain of Alzheimer’s disease patients and associated with risk of AD disease [53, 54, 55, 56].

Protein-protein docking methods Zdock and Rosetta were applied on these proteins to build complex structures between CRBN and the target proteins. The starting structures for CRBN was obtained from the ternary complex structure of CRBN-PROTAC-BRD4 (PDB id: 6BOY) and the ligand mimic E3 ligase moiety was from CRBN-ligand complex structure (PDB id: 4tz4, ligand: Lenalidomide (LVY)). After the complex structures were obtained, Autodock and the protein-ligand docking protocol in Rosetta were performed to generate docking conformations for the selected PROTAC, as shown in the method. 10 docking conformations were generated according to their docking scores and the one closest to the original LVY was selected to perform further interaction analyse. 3D and 2D interactions between the docked PROTACs and the CRBN-target protein complexes were shown in Fig 6.

**Fig 6.**
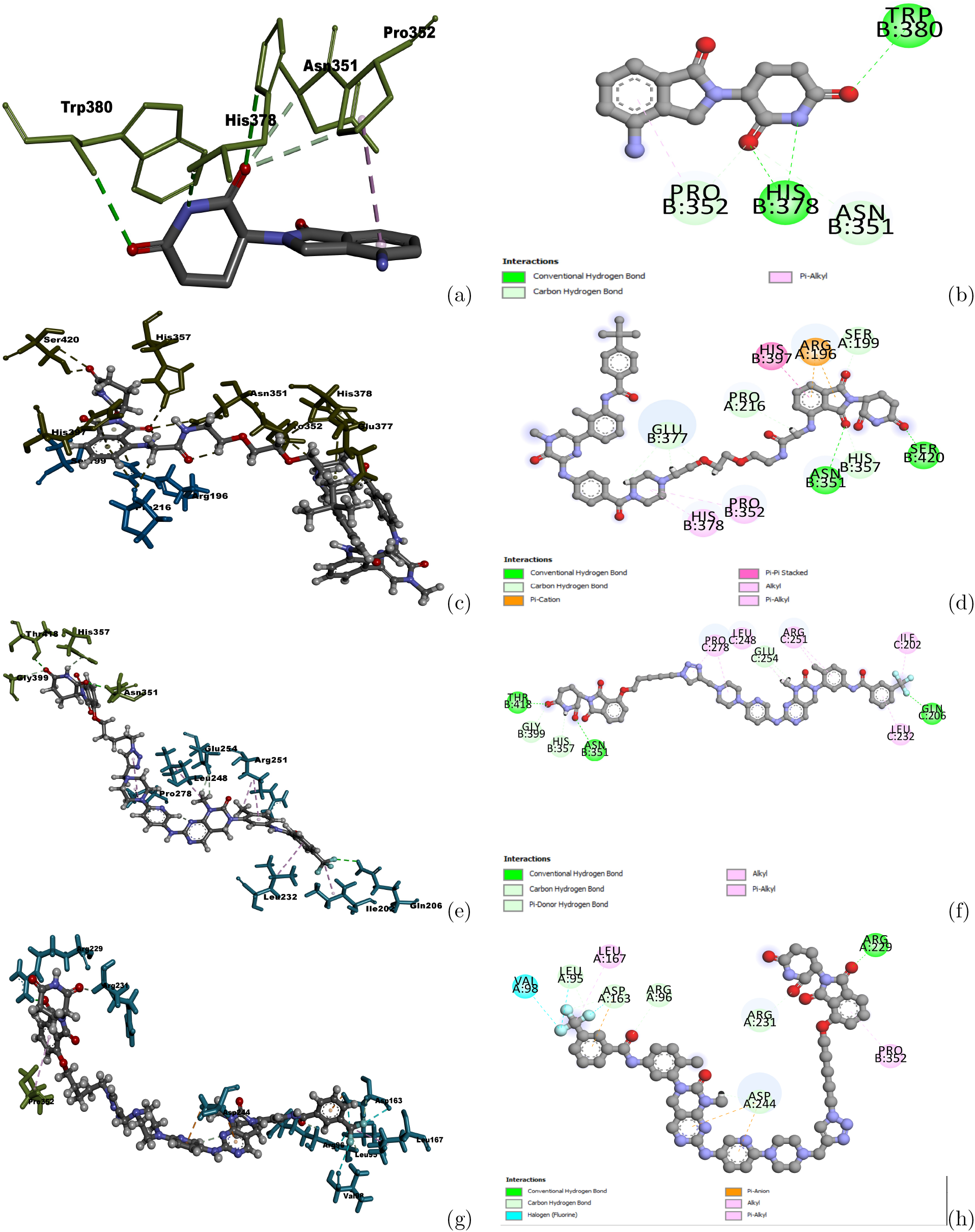
Interactions between predicted PROTACs and the CRBN-target protein complex structures. (a) Interactions between CRBN and its crystallized ligand LVY. (b) 2D interactions between CRBN and its crystallized ligand LVY. (c) Interactions between Protac1222 and the complex structure of P12575 and CRBN. (d) 2D interactions between Protac1222 and the complex structure of P12575 and CRBN. (e) Interactions between Protac2145 and the complex structure of Q9UER7 and CRBN. (f) 2D interactions between Protac2145 and the complex structure of Q9UER7 and CRBN. (g) Interactions between Protac2145 and the complex structure of Q9H1I8 and CRBN. (h) 2D interactions between Protac2145 and the complex structure of Q9H1I8 and CRBN. Dark green sticks represent interacting residues on CRBN, Cyan sticks represent interacting residues on the target protein. Balls and sticks represent the ligand

For P12757, most of the residues on CRBN interacting with the CRBN ligand moiety remained between CRBN and PROTAC1222, including ASN315, HIS378 and PRO352. TRP380 did not interact with CRBN ligand moiety. Instead, SER420 formed a hydrogen bond with the oxygen atom on the ring. For the other two target proteins, there were also several interactions observed between CRBN and the CRBN ligand moiety on the predicted RPOTACs. For example, the interaction between ASN351 and the CRBN ligand moiety for Q9UER7 and the interaction between PRO352 and CRBN ligand moiety for Q9H1I8 remained the same as between CRBN and the co-crystallized ligand LVY. Other interacting residues on these target protein-CRBN complex structures were different with those in the original CRBN-ligand complex structure, but they formed similarly hydrogen bonds with oxygen atoms on the ring of the CRBN ligand moiety. Many residues on these target proteins also formed favorable interactions with the warhead parts on the PROTACs. Interactions between CRBN and CRBN ligand moiety, and between the target proteins and warheads indicate the predicted PROTACs could form a dual binding with CRBN and target proteins and have the potential to bring the target proteins to CRBN and induce the degradation of these proteins.

Key positions were also identified for these three proteins by eSHAP analysis. In order to compare with the docking results, the top ranked positions were mapped on the modeled complex structures for P12757, Q9UER7 and Q9H1I8 and shown in Fig 7. Majority of these key positions were found around their PROTAC binding pockets on the modeled structures. An interesting thing for Q9H1I8 is that the predicted PROTAC binding pockets are also the interface between the human activating signal co-integrator complex ASCC2 and ASCC3 [57]. Leu91, Tyr94, Pro99, Ile 201, Leu256 and Phe270 are all around this PROTAC binding pockets and the interface between ASCC3 and ASCC2. Identified key positions on these predicted model structures showed that our model can provide useful clues about binding residues when designing PROTACs for understudied proteins.

**Fig 7.**
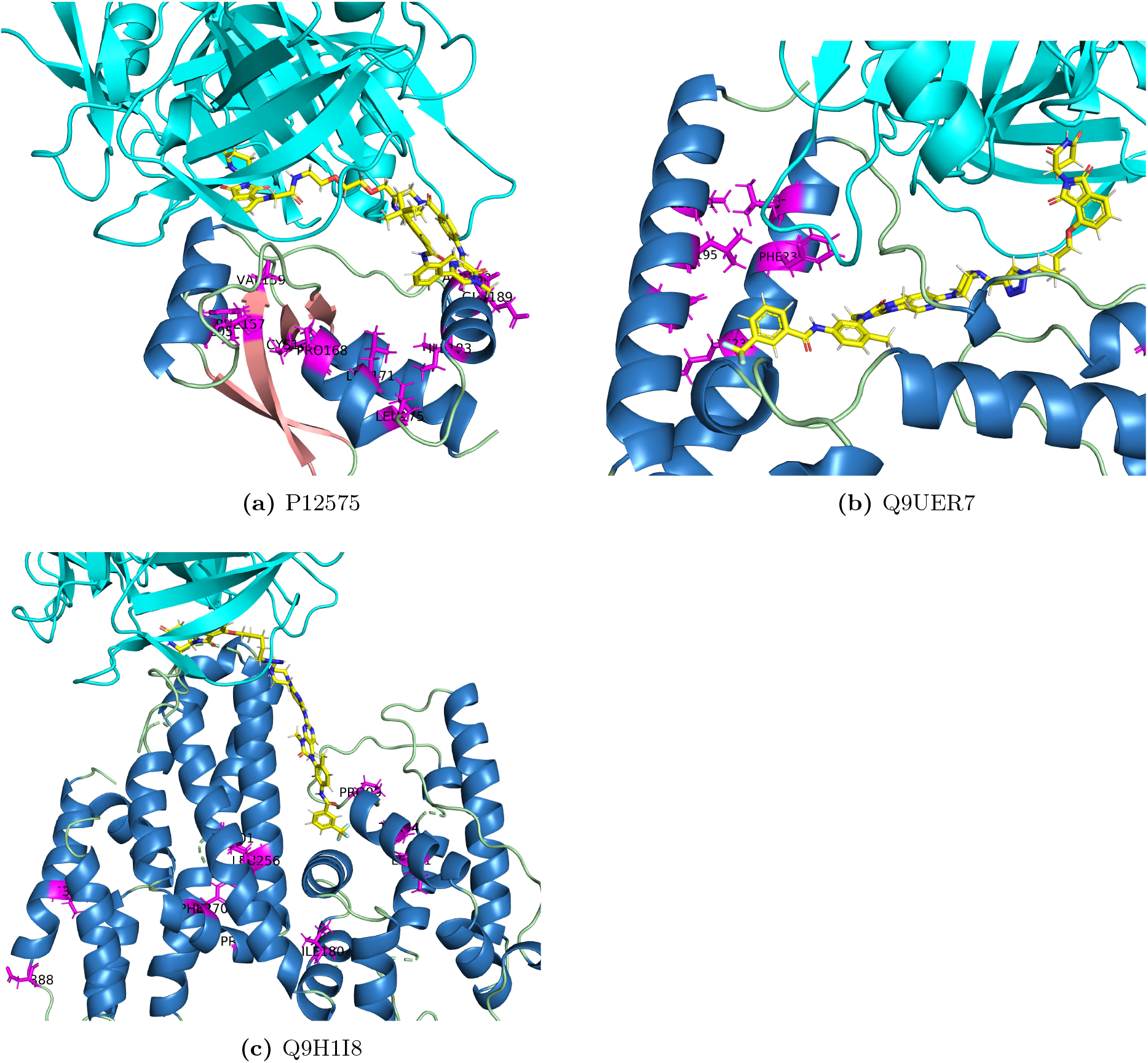
Structural mapping of the key positions for P12575, Q9UER7 and Q9H1I8. Pink represents strands, blue represents helices, green represents loop regions, yellow represents co-crystalized ligand and magenta represents the top ranked positions.

## 3 Conclusion

Protein degradation is a normal process of protein turnover within the cell. It provides a mechanism of quality control during protein folding, an ability to rapidly respond to changing cellular signals, and a mechanism to modulate the pool of available amino acids. The majority of proteins will undergo degradation through the ubiquitin–proteasome system. The ubiquitin E3 ligase family is the largest family in ubiquitin signaling, which includes about 700 members identified or predicted to possess ligase activities [58]. There are three subfamilies of E3 ubiquitin ligases: RING [59], HECT[60], and RING-Between-RING (RBR) E3 ligases [61], which include PARKIN and ARIH1, and mechanistically behave as hybrids between RING and HECT. Currently, most reported PROTACs rely on two E3 ligases, CRBN and VHL. Our training data also depends on the degradation data for CRBN E3 ligase. It remains unknown if the model developed from CRBN can be applied to other E3 ligase. Studies on the protein kinase degradation showed that there are overlaps between CRBN-caused degradation and VHL-caused degradation, which indicates the targets degradable for one E3 ligase could also be potential targets for another E3 ligase.

Currently, the majority of PROTACs are designed from known potent target binding ligands and usually target on well-studied proteins. Many understudied proteins that are potential drug targets have not been explored by the PROTAC. The most promising advantage for the PROTAC should be the ability to induce these proteins for degradation. Our results for understudied human proteins could be a starting point to develop PROTACs for these proteins. With our prediction results, more detailed studies like sequence and structure comparison, complex structure modeling and linker modeling can be applied and help to design PROTACs for these proteins.

Our method is the first machine learning method which can use the features from protein sequence to predict PROTAC-induced degradation for target proteins. In this work, 23 different feature descriptors, three classification methods and combination of them were used to build machine learning models. These models were trained and validated by protein kinase degradation data and tested on other degradable proteins from ProtacDB. A final model was built based on all degradation data from both the training set and test set. In this way, a throughout examination about protein sequence features, classification methods were performed on currently available PROTAC-induced degradation data and the final model obtained here can be used to do Pre-PROTAC screening to get possible target proteins which can be degraded by CRBN. Compared with other computational methods, this method can screen a large number of proteins in a short time. With more and more degradation data available in the future, accuracy of such model could be largely improved.

eSHAP analysis and key position identification on protein kinases and BRD4 showed that our model can be able to predict key positions for PROTAC-induced degradation. Limited by the current availability of degradation data, there are still many noises for such prediction. Combined with other structural features and knowledge about the target proteins, more accurate prediction could be expected and would provide useful information to help design PRATACs for the target proteins.

The number of data points is the main limitation in this work. But our model is not just trying to differentiate which structural or sequential features that might lead to protein kinases susceptible to targeted protein degradation. In the training set, there are 211 positive targets and 234 negative targets and in the test set, there are 62 other proteins. PrePROTAC also achieved reasonable accuracy for the proteons in the test set, as shown in Fig 3. And our final model put all the data points together, including not only protein kinases but also other proteins. It is hard to predict all degradable proteins based on currently available data. Even if only a small number of the degradable proteins could be found with our model, it will also be a good start for PROTACs design. The performance of our model can be further improved when there are more degradation data available in the future.

In our current work, only sequence information is considered to build and train the models. Structural features of the binding pockets for both E3 ligases and the target proteins would be able to provide additional information and guide the Pre-PROTAC prediction. Current protein structure prediction methods, like alpha-fold [62], can build model structures for proteins without known crystal structures. However, these model structures need to be selected carefully before being used to extract information since there are many randomly built loops in the model structures. Models including reliable structural features will be able to improve performance for Pre-PROTAC prediction.

## 4 Method

Random forest (RF) and gradient boosting tree (GBT) classification methods implemented in scikit-learn [63] were evaluated in this work and the one with the best performance was selected to predict whether a protein could be degraded by PROTAC-induced protein degradation. These models were trained based on human kinases degradation data from Fischer’s group [64] and tested on other degradation data from PROTAC-DB [65]. Pre-trained features from iFeature [40], D-Script [41], and ESM [42] were selected as the input for the RF and GBT classification methods. Combination of different classification methods and features were tested by the cross validation method on the training set and the RF classification with ESM features had the best performance. The RF and GBT classfication with ESM features were further examined on the test set. At the end, the training set and the test set was put together to build the final PrePROTAC model.

### 4.1 Training data set

One of the difficulties on PROTAC-induced protein degradation is few data available for systematic modeling studies. Comparing with traditional drug discovery, only a small number of PROTACs and targeted proteins are reported. Recently, Fischer’s group developed a large library of kinase-targeting degraders and identified degradable protein-kinases among human protein kinases and kinase-like proteins. [64]. Based on their results, 445 protein kinases were selected as the training set, including 201 proteins degraded by CRBN-recruiting degraders, 95 proteins degraded by VHL-recruiting degraders and 234 proteins which cannot be degraded by any degraders. There are some overlaps between proteins degraded by CRBN and VHL. For example, among the 95 proteins degraded by VHL, only 10 of them are exclusively degraded by VHL-recruiting degraders. Distributions of CRBN and VHL induced degradations in different protein kinase families was shown in S24 Fig. Labels for the proteins in the training set are based on the experimental results. Since the majority degradation are recruited by CRBN, the machine learning models should be trained for CRBN targeted degradation. In the training set, if a target is degraded by CRBN, the target is labeled as positive, otherwise, negative.

### 4.2 Test data set

In order to evaluate the machine learning classification model trained by the CRBN-protein kinase pairs, additional PROTAC-induced degradation data was collected from PROTAC-DB [65] to build an external test data set. PROTAC-DB is an online database which gather information of PROTACs by searching PubMed with keywords of ‘degrader* or PROTAC or proteolysis targeting chimera’. 62 proteins were found to be degraded by CRBN or VHL-recruiting degraders and not shown in protein kinase training set. Among them, 8 proteins are VHL targets and have no evidence to confirm whether they can be degraded by CRBN or not. The other 54 proteins were used as positive samples in the test data set. Till now, known negative degradation data are very few. Thus, in order to get a negative control, we randomly selected representative proteins in different super families but in the same SCOP [66, 67] fold with the 54 degradable proteins. These proteins are real proteins and have similar fold structures with the degradable proteins. There are some chances for these proteins in the SCOP fold set to be degraded by CRBN. Here, they were used as negative controls and may result in a lower performance for the classification model when some of them are actually degradable.

### 4.3 Machine learning methods

The goal of this study is to identify which proteins can be induced by PROTAC molecules and further degraded by CRBN. Two different classification methods were investigated for this purpose, including RF and GBT classification methods. RF is one of the most commonly used classification algorithm, which consists of many decisions trees. The fundamental concept, “the wisdom of crowds”, make it simple but powerful. GBT method optimize a cost function over function space and combine weak learners into a single strong learner in an iterative way.

Here, these two different classification methods were trained to learn from the features describing proteins and distinguish between degradable and undegradable proteins in the training set. Four Hyper-parameters, n_estimators, max_depth, min_samples_split and min_samples_leaf were tuned through 5-fold grid-search cross validation method and the ones with the best ROC-AUC scores were chosen for the next step modeling.

### 4.4 Pretrained features

Since there are only several hundreds data points for PROTAC induced degradation, it is not practical to perform large scale training. Here, Pre-trained features from other large scale protein language modeling were used as an initial input for the random forest classification method, including sequence features from iFeature [40], pre-trained protein-protein interaction contact features from D-SCRIPT [41], and pre-trained protein sequence features from ESM [42]. These features are extracted from a large number of protein sequences or structures and will help our classification models to capture useful information when being used as input for these models.

#### 4.4.1 Sequence features from iFeature

We first tested the features based on structural and physio-chemical descriptors extracted from protein sequence data. iFeature [40] is a python-based toolkit which integrates 18 major sequence encoding schemes and 53 different types of feature descriptors. The number of features for these descriptors range from hundreds to thousands. In order to avoid over-fitting, 21 descriptors of them which contain less than 1000 features were selected as the features for the classification models. As shown in table S1, these descriptors can be separated into six groups, including amino acid composition related [68, 69, 70, 71], grouped amino acid composition related, distribution of amino acid properties related [72] from AAindex database [73], amino acid distribution patterns related [74], conjoint triad descriptor related [75] and sequence-order feature related groups [76].

#### 4.4.2 Contact features from D-SCRIPT

In PROTAC induced degradation, targeted proteins and E3 ligase will form complex structures. But such complex structures are different with the naturally formed protein-protein complex since PROTAC binds to the interface between them and hold them to stay together. That is why current protein-protein interaction prediction methods cannot directly predict such pairs. However, the information hidden in the protein-protein interaction is still useful for the prediction of PROTAC induced degradation because the formation of E3 ligase-target protein complex structures still follows basic physical-chemical principles. Thus, the pretrained features used in the protein-protein interaction prediction method can be a start point for PROTAC induced degradation.

In this work, the contact feature in D-SCRIPT [41], a deep learning method for predicting a physical interaction between two proteins given just their sequences, was applied. Starting from the pre-trained embedding models E1 and E2 which capture both local and global structural information from both sequences in one CRBN-target pair, a projection module was used to reduce the dimensions of E1 and E2 by using a fully-connected linear layer. A residue contact module was then used to model the interactions between the residues of each protein. This step generated a *n*-by-*m* contact prediction matrix which indicates the predicted probability that two residues (one in CRBN and the other one in target protein) are in contact. Here, *n* represents the sequence length of the CRBN and *m* represents the length of the target protein. This matrix was used as the initial features in our model. In order to get a fix-length array for each pair, a max pooling operation was used to change the matrix to *n*-dimensional array. This array represents *n*-dimensional contact features for each sample in the random forest classification.

#### 4.4.3 Features from ESM-1b transformer

Recently, Facebook AI research group developed a deep contextual language model with unsupervised learning to train on 86 billion amino acids across 250 million protein sequences [42]. The resulting model contains information about biological properties in its representations. The representations are learned from sequence data alone, and ESM-1b model outperforms all tested single-sequence protein language models across a range of structure prediction tasks. Here, we used the output embedding features from ESM-1b model as the input features to train the random forest classification model to identity PROTAC-induced degradable proteins.

### 4.5 Performance evaluation

Performance of RF and GBT models with different features were evaluated by the repeated Stratified 5-Fold cross validation method. Average area under the curve of Receiver operating characteristic curve (ROC-AUC) and precision score were calculated and compared between these models. The model with the highest score was selected for further prediction.

### 4.6 Soft voting model with ESM feature

In order to use all the PROTAC-induced degradation data, proteins in the training set and external test set were combined and a final model was trained for the whole data set. By using repeated Stratified K-Fold cross validation with n_splits as 5 and n_repeats as 2, ten groups of randomly selected training set and validation set were generated and each set contained approximately the same percentage of positive and negative samples as in the whole data set. Ten random forest classification models were trained and validated on these data sets. Then a soft voting classifying model was built by summing the predicted probabilities from the ten models with equal weight. Based on the correlation between average false positive rate and threshold from ten validation sets, if the threshold is set as 0.85, the false positive rate will be 0.004, which means only one of the negative sample is predicted as positive since there are 241 negative samples in the data set. If the threshold is set as 0.90, the false positive rate will be 0 which means all of the negative samples will be predicted as negative. In order to select the potential degradable proteins with high confidence, the threshold of 0.90 will be used to select positive hits in the prediction. This model could be used before PROTACs design to select target proteins and is called PrePROTAC model.

### 4.7 eSHAP analysis of PrePROTAC model identifies key residues contributing to PROTAC activities

SHAP (SHapley Additive exPlanations)[77] [78] [79] values are widely used to explain machine learning models. TreeExplainer in the shap python package was applied to get the SHAP values for the features used in the PrePROTAC model. The 20 embedding features with the highest SHAP values were selected and these features should play important roles in the prediction of our model. Since these features are embedding attributes, their specific meaning is unknown. In order to investigate which residues on the protein make key contributions to PROTAC activity, an in silico mutagenesis was performed on the proteins in the training set. Each residue was mutated sequentially once a time to the amino acid with the opposite property. For example positive charged amino acids mutated to negative charged amino acids and polar amino acids mutated to hydrophobic amino acids. For each position on the protein sequence, the differences of the 20 selected embedding features between the mutated sequences and the original sequence was calculated according to the following formula:

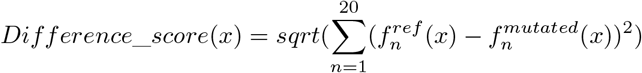

The importance of all the amino acid positions were measured by the difference scores. This embedding SHAP (eSHAP) analysis could help identify the residues which contribute most on the selected features. The bigger the difference score is, the higher impact of this position has on these features. The positions with the top-ranked difference scores were selected as the key positions for this protein.

### 4.8 Protein-protein docking and protein-ligand docking

In order to design potential PROTACs for the target proteins, protein-protein docking and protein-ligand docking methods were applied on target proteins to build complex structures between E3-ligases and target proteins, and to predict PROTACs which were able to interact with these target proteins. Zdock [80] was firstly performed to bring CRBN and target proteins together. The interface residues of CRBN were used to guide the protein-protein docking in Zdock. A local protein-protein docking protocol in Rosetta [81] was used to refine the complex structure obtained from Zdock. Then, the CRBN-target protein complex structure was decided by choosing the complex structure with the lowest energy. Autodock [82] was then applied to screen the 275 warheads in ProtacDB and select the warheads which could dock into the binding pocket for the target protein. Warheads with lower docking scores were selected and their matched PROTACs were chosen from ProtacDB. Then these PROTACs were docked into the predicted complex structures by Rosetta protein-ligand docking to generate docking conformations with lower energies.

## Author Contributions

Li Xie prepared data, implemented the algorithms, performed the experiments, and wrote the manuscript; Lei Xie conceived and planned the experiments, wrote the manuscript.

## Data and software availability

All the data and source code for the model training and prediction can be found at https://github.com/XieResearchGroup/PrePROTAC.git

## Supporting information

**S1 Table. Optimized hyper-parameters for the random forest classification models with different features.**

**S2 Table. Optimized hyper-parameters for the gradient boosting classification models with different features.**

**S3 Table. Protein information and probability scores of 615 human proteins predicted to be targets of PROTAC.**

**S1 Fig. Average ROC-AUC scores for random forest and gradient boosting classification models with different features.**

**S2 Fig. ROC-AUC curves for random forest classification models with different features.**

**S3 Fig. False positive rate - threshold curves for random forest classification models with different features.**

**S4 Fig. Precision - recall curves for random forest classification models with different features.**

**S5 Fig. ROC-AUC curves for the gradient boosting classification models with different features.**

**S6 Fig. False positive rate - threshold curves for the gradient boosting classification models with different features.**

**S7 Fig. Precision - recall curves for the gradient boosting classification models with different features.**

**S8 Fig. Average ROC-AUC scores and precision scores for random forest models with combined features.**

**S9 Fig. ROC-AUC, false positive rate-threshold, precision-recall curves for the soft voting model and consensus model on the training set.**

**S10 Fig. ROC-AUC, false positive rate-threshold, precision-recall curves for the soft voting model and consensus model on the test set with SCOP proteins as the negative samples.**

**S11 Fig. Structural mapping of the top ranked positions for positive and negative samples in protein kinase subgroup 1 and multiple sequence alignments of the top ranked positions in this subgroup.**

**S12 Fig. Structural mapping of the top ranked positions for positive and negative samples in protein kinase subgroup 2 and multiple sequence alignments of the top ranked positions in this subgroup.**

**S13 Fig. Structural mapping of the top ranked positions for positive and negative samples in protein kinase subgroup 3 and multiple sequence alignments of the top ranked positions in this subgroup.**

**S14 Fig. Structural mapping of the top ranked positions for positive and negative samples in protein kinase subgroup 4 and multiple sequence alignments of the top ranked positions in this subgroup.**

**S15 Fig. Structural mapping of the top ranked positions for positive and negative samples in protein kinase subgroup 5 and multiple sequence alignments of the top ranked positions in this subgroup.**

**S16 Fig. Structural mapping of the top ranked positions for positive and negative samples in protein kinase subgroup 6 and multiple sequence alignments of the top ranked positions in this subgroup.**

**S17 Fig. Structural mapping of the top ranked positions for positive and negative samples in protein kinase subgroup 7 and multiple sequence alignments of the top ranked positions in this subgroup.**

**S18 Fig. Structural mapping of the top ranked positions for positive and negative samples in protein kinase subgroup 8 and multiple sequence alignments of the top ranked positions in this subgroup.**

**S19 Fig. Structural mapping of the top ranked positions for positive and negative samples in protein kinase subgroup 9 and multiple sequence alignments of the top ranked positions in this subgroup.**

**S20 Fig. Structural mapping of the top ranked positions for positive and negative samples in protein kinase subgroup 10 and multiple sequence alignments of the top ranked positions in this subgroup.**

**S21 Fig. Structural mapping of the top ranked positions for positive and negative samples in protein kinase subgroup 11 and multiple sequence alignments of the top ranked positions in this subgroup.**

**S22 Fig. Structural mapping of the top ranked positions for positive and negative samples in protein kinase subgroup 12 and multiple sequence alignments of the top ranked positions in this subgroup.**

**S23 Fig. Distribution of predicted probability scores for the whole human proteome.**

**S24 Fig. Distributions of CRBN induced degradations in different protein kinase families.**

## Acknowledgement

This work has been supported by the National Institute of General Medical Sciences of National Institute of Health (R01GM122845) and the National Institute on Aging of the National Institute of Health (R01AD057555).

## References

[1] Hopkins AL, Groom CR. The druggable genome. Nature Reviews Drug Discovery. 2002;1(9):727–730.

[2] Overington JP, Al-Lazikani B, Hopkins AL. How many drug targets are there? Nature Reviews Drug Discovery. 2006;5(12):993–996.

[3] Lazo JS, Sharlow ER. Drugging Undruggable Molecular Cancer Targets. Annual Review of Pharmacology and Toxicology. 2016;56:23–40.

[4] Nalawansha DA, Crews CM. PROTACs: An Emerging Therapeutic Modality in Precision Medicine. Cell Chemical Biology. 2020;27(8):998–1014.

[5] Paiva SL, Crews CM. Targeted protein degradation: elements of PROTAC design. Current Opinion in Chemical Biology. 2019;50:111–119. doi:10.1016/j.cbpa.2019.02.022.

[6] Smith BE, Wang SL, Jaime-Figueroa S, Harbin A, Wang J, Hamman BD, et al. Differential PROTAC substrate specificity dictated by orientation of recruited E3 ligase. Nature Communications. 2019;10(1). doi:10.1038/s41467-018-08027-7.

[7] Gechijian LN, Buckley DL, Lawlor MA, Reyes JM, Paulk J, Ott CJ, et al. Functional TRIM24 degrader via conjugation of ineffectual bromodomain and VHL ligands. Nature Chemical Biology. 2018;14(4):405–412.

[8] Bassi ZI, Fillmore MC, Miah AH, Chapman TD, Maller C, Roberts EJ, et al. Modulating PCAF/GCN5 Immune Cell Function through a PROTAC Approach. ACS Chemical Biology. 2018;13(10):2862–2867. doi:10.1021/acschembio.8b00705.

[9] Cromm PM, Samarasinghe KTG, Hines J, Crews CM. Addressing Kinase-Independent Functions of Fak via PROTAC-Mediated Degradation. Journal of the American Chemical Society. 2018;140(49):17019–17026. doi:10.1021/jacs.8b08008.

[10] Degorce SL, Tavana O, Banks E, Crafter C, Gingipalli L, Kouvchinov D, et al. Discovery of Proteolysis-Targeting Chimera Molecules that Selectively Degrade the IRAK3 Pseudokinase. Journal of Medicinal Chemistry. 2020;63(18):10460–10473. doi:10.1021/acs.jmedchem.0c01125.

[11] Crews CM. Targeting the Undruggable Proteome: The Small Molecules of My Dreams. Chemistry & Biology. 2010;17(6):551–555. doi:10.1016/j.chembiol.2010.05.011.

[12] Schapira M, Calabrese MF, Bullock AN, Crews CM. Targeted protein degradation: expanding the toolbox. Nature Reviews Drug Discovery. 2019;18(12):949–963. doi:10.1038/s41573-019-0047-y.

[13] Lai AC, Crews CM. Induced protein degradation: an emerging drug discovery paradigm. Nature Reviews Drug Discovery. 2016;16(2):101–114. doi:10.1038/nrd.2016.211.

[14] Bondeson DP, Smith BE, Burslem GM, Buhimschi AD, Hines J, Jaime-Figueroa S, et al. Lessons in PROTAC Design from Selective Degradation with a Promiscuous Warhead. Cell Chemical Biology. 2018;25(1):78–87.e5. doi:10.1016/j.chembiol.2017.09.010.

[15] Gechijian LN, Buckley DL, Lawlor MA, Reyes JM, Paulk J, Ott CJ, et al. Functional TRIM24 degrader via conjugation of ineffectual bromodomain and VHL ligands. Nature Chemical Biology. 2018;14(4):405–412. doi:10.1038/s41589-018-0010-y.

[16] Burslem GM, Smith BE, Lai AC, Jaime-Figueroa S, McQuaid DC, Bondeson DP, et al. The Advantages of Targeted Protein Degradation Over Inhibition: An RTK Case Study. Cell Chemical Biology. 2018;25(1):67–77.e3. doi:10.1016/j.chembiol.2017.09.009.

[17] Cromm PM, Samarasinghe KTG, Hines J, Crews CM. Addressing Kinase-Independent Functions of Fak via PROTAC-Mediated Degradation. Journal of the American Chemical Society. 2018;140(49):17019–17026. doi:10.1021/jacs.8b08008.

[18] Popow J, Arnhof H, Bader G, Berger H, Ciulli A, Covini D, et al. Highly Selective PTK2 Proteolysis Targeting Chimeras to Probe Focal Adhesion Kinase Scaffolding Functions. Journal of Medicinal Chemistry. 2019;62(5):2508–2520. doi:10.1021/acs.jmedchem.8b01826.

[19] Vogelmann A, Robaa D, Sippl W, Jung M. Proteolysis targeting chimeras (PROTACs) for epigenetics research. Current Opinion in Chemical Biology. 2020;57:8–16. doi:10.1016/j.cbpa.2020.01.010.

[20] Winter GE, Buckley DL, Paulk J, Roberts JM, Souza A, Dhe-Paganon S, et al. Phthalimide conjugation as a strategy for in vivo target protein degradation. Science. 2015;348(6241):1376–1381. doi:10.1126/science.aab1433.

[21] Gadd MS, Testa A, Lucas X, Chan KH, Chen W, Lamont DJ, et al. Structural basis of PROTAC cooperative recognition for selective protein degradation. Nature Chemical Biology. 2017;13(5):514–521. doi:10.1038/nchembio.2329.

[22] Raina K, Lu J, Qian Y, Altieri M, Gordon D, Rossi AMK, et al. PROTAC-induced BET protein degradation as a therapy for castration-resistant prostate cancer. Proceedings of the National Academy of Sciences. 2016;113(26):7124–7129. doi:10.1073/pnas.1521738113.

[23] Schiedel M, Herp D, Hammelmann S, Swyter S, Lehotzky A, Robaa D, et al. Chemically Induced Degradation of Sirtuin 2 (Sirt2) by a Proteolysis Targeting Chimera (PROTAC) Based on Sirtuin Rearranging Ligands (SirReals). Journal of Medicinal Chemistry. 2017;61(2):482–491. doi:10.1021/acs.jmedchem.6b01872.

[24] An Z, Lv W, Su S, Wu W, Rao Y. Developing potent PROTACs tools for selective degradation of HDAC6 protein. Protein & Cell. 2019;10(8):606–609. doi:10.1007/s13238-018-0602-z.

[25] Smalley JP, Adams GE, Millard CJ, Song Y, Norris JKS, Schwabe JWR, et al. PROTAC-mediated degradation of class I histone deacetylase enzymes in corepressor complexes. Chemical Communications. 2020;56(32):4476–4479. doi:10.1039/d0cc01485k.

[26] Dong H, Liu S, Zhang X, Chen S, Kang L, Chen Y, et al. An Allosteric PRC2 Inhibitor Targeting EED Suppresses Tumor Progression by Modulating the Immune Response. Cancer Research. 2019;79(21):5587–5596. doi:10.1158/0008-5472.can-19-0428.

[27] Hsu JHR, Rasmusson T, Robinson J, Pachl F, Read J, Kawatkar S, et al. EED-Targeted PROTACs Degrade EED, EZH2, and SUZ12 in the PRC2 Complex. Cell Chemical Biology. 2020;27(1):41–46.e17. doi:10.1016/j.chembiol.2019.11.004.

[28] Potjewyd F, Turner AMW, Beri J, Rectenwald JM, Norris-Drouin JL, Cholensky SH, et al. Degradation of Polycomb Repressive Complex 2 with an EED-Targeted Bivalent Chemical Degrader. Cell Chemical Biology. 2020;27(1):47–56.e15. doi:10.1016/j.chembiol.2019.11.006.

[29] Salami J, Alabi S, Willard RR, Vitale NJ, Wang J, Dong H, et al. Androgen receptor degradation by the proteolysis-targeting chimera ARCC-4 outperforms enzalutamide in cellular models of prostate cancer drug resistance. Communications Biology. 2018;1(1). doi:10.1038/s42003-018-0105-8.

[30] Buhimschi AD, Armstrong HA, Toure M, Jaime-Figueroa S, Chen TL, Lehman AM, et al. Targeting the C481S Ibrutinib-Resistance Mutation in Bruton’s Tyrosine Kinase Using PROTAC-Mediated Degradation. Biochemistry. 2018;57(26):3564–3575. doi:10.1021/acs.biochem.8b00391.

[31] Mares A, Miah AH, Smith IED, Rackham M, Thawani AR, Cryan J, et al. Extended pharmacodynamic responses observed upon PROTAC-mediated degradation of RIPK2. Communications Biology. 2020;3(1). doi:10.1038/s42003-020-0868-6.

[32] Zaidman D, Prilusky J, London N. PRosettaC: Rosetta Based Modeling of PROTAC Mediated Ternary Complexes. Journal of Chemical Information and Modeling. 2020;60(10):4894–4903. doi:10.1021/acs.jcim.0c00589.

[33] Bai N, Miller SA, Andrianov GV, Yates M, Kirubakaran P, Karanicolas J. Rationalizing PROTAC-Mediated Ternary Complex Formation Using Rosetta. Journal of Chemical Information and Modeling. 2021;61(3):1368–1382. doi:10.1021/acs.jcim.0c01451.

[34] Nowak RP, DeAngelo SL, Buckley D, He Z, Donovan KA, An J, et al. Plasticity in binding confers selectivity in ligand-induced protein degradation. Nature Chemical Biology. 2018;14(7):706–714. doi:10.1038/s41589-018-0055-y.

[35] Drummond ML, Williams CI. In Silico Modeling of PROTAC-Mediated Ternary Complexes: Validation and Application. Journal of Chemical Information and Modeling. 2019;59(4):1634–1644. doi:10.1021/acs.jcim.8b00872.

[36] Drummond ML, Henry A, Li H, Williams CI. Improved Accuracy for Modeling PROTAC-Mediated Ternary Complex Formation and Targeted Protein Degradation via New In Silico Methodologies. Journal of Chemical Information and Modeling. 2020;60(10):5234–5254. doi:10.1021/acs.jcim.0c00897.

[37] Lebraud H, Wright DJ, Johnson CN, Heightman TD. Protein Degradation by In-Cell Self-Assembly of Proteolysis Targeting Chimeras. ACS Central Science. 2016;2(12):927–934. doi:10.1021/acscentsci.6b00280.

[38] Testa A, Hughes SJ, Lucas X, Wright JE, Ciulli A. Structure-Based Design of a Macrocyclic PROTAC. Angewandte Chemie International Edition. 2019;59(4):1727–1734. doi:10.1002/anie.201914396.

[39] Imrie F, Bradley AR, van der Schaar M, Deane CM. Deep Generative Models for 3D Linker Design. Journal of Chemical Information and Modeling. 2020;60(4):1983–1995. doi:10.1021/acs.jcim.9b01120.

[40] Chen Z, Zhao P, Li F, Leier A, Marquez-Lago TT, Wang Y, et al. iFeature: a Python package and web server for features extraction and selection from protein and peptide sequences. Bioinformatics. 2018;34(14):2499–2502.

[41] Sledzieski S, Singh R, Cowen L, Berger B. Sequence-based prediction of protein-protein interactions: a structure-aware interpretable deep learning model. biorxiv. 2021;.

[42] Rives A, Meier J, Sercu T, Goyal S, Lin Z, Liu J, et al. Biological structure and function emerge from scaling unsupervised learning to 250 million protein sequences. Proceedings of the National Academy of Sciences of the United States of America. 2021;118(15):e2016239118.

[43] Modi V, Dunbrack RL. A structurally-validated multiple sequence alignment of 497 human protein kinase domains. Scientific reports. 2019;9(1):1–16.

[44] Kumar S, Stecher G, Li M, Knyaz C, Tamura K. MEGA X: molecular evolutionary genetics analysis across computing platforms. Molecular biology and evolution. 2018;35(6):1547.

[45] Letunic I, Bork P. Interactive tree of life (iTOL) v3: an online tool for the display and annotation of phylogenetic and other trees. Nucleic acids research. 2016;44(W1):W242–W245.

[46] Hanks SK, Quinn AM, Hunter T. The protein kinase family: conserved features and deduced phylogeny of the catalytic domains. Science. 1988;241(4861):42–52.

[47] Manning G, Whyte DB, Martinez R, Hunter T, Sudarsanam S. The protein kinase complement of the human genome. Science. 2002;298(5600):1912–1934.

[48] Lu J, Qian Y, Altieri M, Dong H, Wang J, Raina K, et al. Hijacking the E3 ubiquitin ligase cereblon to efficiently target BRD4. Chemistry & biology. 2015;22(6):755–763.

[49] Finan C, Gaulton A, Kruger FA, Lumbers RT, Shah T, Engmann J, et al. The druggable genome and support for target identification and validation in drug development. Science translational medicine. 2017;9(383).

[50] Sheils TK, Mathias SL, Kelleher KJ, Siramshetty VB, Nguyen DT, Bologa CG, et al. TCRD and Pharos 2021: mining the human proteome for disease biology. Nucleic Acids Research. 2020;49(D1):D1334–D1346.

[51] Finan C, Gaulton A, Kruger FA, Lumbers RT, Shah T, Engmann J, et al. The druggable genome and support for target identification and validation in drug development. Science Translational Medicine. 2017;9:eaag1166.

[52] Piñero J, Ramírez-Anguita JM, Saüch-Pitarch J, Ronzano F, Centeno E, Sanz F, et al. The DisGeNET knowledge platform for disease genomics: 2019 update. Nucleic Acids Research. 2020;48(D1):D845–D855.

[53] Qu J, Nakamura T, Cao G, Holland EA, McKercher SR, Lipton SA. S-Nitrosylation activates Cdk5 and contributes to synaptic spine loss induced by *β*-amyloid peptide. Proceedings of the National Academy of Sciences. 2011;108(34):14330–14335.

[54] Haun F, Nakamura T, Shiu AD, Cho DH, Tsunemi T, Holland EA, et al. S-nitrosylation of dynamin-related protein 1 mediates mutant huntingtin-induced mitochondrial fragmentation and neuronal injury in Huntington’s disease. Antioxidants & redox signaling. 2013;19(11):1173–1184.

[55] Walter S, Atzmon G, Demerath EW, Garcia ME, Kaplan RC, Kumari M, et al. A genome-wide association study of aging. Neurobiology of aging. 2011;32(11):2109–e15.

[56] Castillo E, Leon J, Mazzei G, Abolhassani N, Haruyama N, Saito T, et al. Comparative profiling of cortical gene expression in Alzheimer’s disease patients and mouse models demonstrates a link between amyloidosis and neuroinflammation. Scientific reports. 2017;7(1):1–16.

[57] Jia J, Absmeier E, Holton N, Pietrzyk-Brzezinska AJ, Hackert P, Bohnsack KE, et al. The interaction of DNA repair factors ASCC2 and ASCC3 is affected by somatic cancer mutations. Nature communications. 2020;11(1):1–13.

[58] Li W, Bengtson MH, Ulbrich A, Matsuda A, Reddy VA, Orth A, et al. Genome-Wide and Functional Annotation of Human E3 Ubiquitin Ligases Identifies MULAN, a Mitochondrial E3 that Regulates the Organelle’s Dynamics and Signaling. PLoS ONE. 2008;3(1):e1487. doi:10.1371/journal.pone.0001487.

[59] Deshaies RJ, Joazeiro CAP. RING Domain E3 Ubiquitin Ligases. Annual Review of Biochemistry. 2009;78(1):399–434. doi:10.1146/annurev.biochem.78.101807.093809.

[60] Berndsen CE, Wolberger C. New insights into ubiquitin E3 ligase mechanism. Nature Structural & Molecular Biology. 2014;21(4):301–307. doi:10.1038/nsmb.2780.

[61] Spratt DE, Walden H, Shaw GS. RBR E3 ubiquitin ligases: new structures, new insights, new questions. Biochemical Journal. 2014;458(3):421–437. doi:10.1042/bj20140006.

[62] Jumper J, Evans R, Pritzel A, Green T, Figurnov M, Ronneberger O, et al. Highly accurate protein structure prediction with AlphaFold. Nature. 2021;596(7873):583–589.

[63] Pedregosa F, Varoquaux G, Gramfort A, Michel V, Thirion B, Grisel O, et al. Scikit-learn: Machine Learning in Python. Journal of Machine Learning Research. 2011;12:2825–2830.

[64] Donovan KA, Ferguson FM, Bushman JW, Eleuteri NA, Bhunia D, Ryu S, et al. Mapping the Degradable Kinome Provides a Resource for Expedited Degrader Development. cell. 2020;183(6):1714–1731.

[65] Weng G, Shen C, Cao D, Gao J, Dong X, He Q, et al. PROTAC-DB: an online database of PROTACs. Nucleic Acids Research. 2021;49(D1):D1381–D1387.

[66] Andreeva A, Howorth D, Chothia C, Kulesha E, Murzin AG. SCOP2 prototype: a new approach to protein structure mining. Nucleic Acids Research. 2013;42(D1):D310–D314. doi:10.1093/nar/gkt1242.

[67] Andreeva A, Kulesha E, Gough J, Murzin AG. The SCOP database in 2020: expanded classification of representative family and superfamily domains of known protein structures. Nucleic Acids Research. 2019;48(D1):D376–D382. doi:10.1093/nar/gkz1064.

[68] Chen X, Qiu JD, Shi SP, Suo SB, Huang SY, Liang RP. Incorporating key position and amino acid residue features to identify general and species-specific Ubiquitin conjugation sites. Bioinformatics. 2013;29(13):1614–1622.

[69] Liu LM, Xu Y, Chou KC. iPGK-PseAAC: Identify Lysine Phosphoglycerylation Sites in Proteins by Incorporating Four Different Tiers of Amino Acid Pairwise Coupling Information into the General PseAAC. Medicinal Chemistry. 2017;13(6):552–559.

[70] Saravanan V, Gautham N. Harnessing Computational Biology for Exact Linear B-Cell Epitope Prediction: A Novel Amino Acid Composition-Based Feature Descriptor. OMICS. 2015;19(10):648–658.

[71] Bhasin M, Raghava GPS. Classification of nuclear receptors based on amino acid composition and dipeptide composition. Journal of Biological Chemistry. 2004;279(22):23262–23266.

[72] Sokal RR, Thomson BA. Population structure inferred by local spatial autocorrelation: an example from an Amerindian tribal population. The American Journal of Physical Anthropology. 2006;129(1):121–131.

[73] Kawashima S, Pokarowski P, Pokarowska M, Kolinski A, Katayama T, Kanehisa M. AAindex: amino acid index database, progress report 2008. Nucleic Acids Research. 2008;36:D202–D205.

[74] Dubchak I, Muchnikt I, Holbrook SR, hou Kim S. Prediction of protein folding class using global description of amino acid sequence. Proceedings of the National Academy of Sciences of the United States of America. 1995;92(19):8700–8704.

[75] Shen J, Zhang J, Luo X, Zhu W, Yu K, Chen K, et al. Predicting protein-protein interactions based only on sequences information. Proceedings of the National Academy of Sciences of the United States of America. 2007;104(11):4337–4341.

[76] Chou KC, Cai YD. Prediction of protein subcellular locations by GO-FunD-PseAA predictor. Biochemical and Biophysical Research Communications. 2004;320(4):1236–1239.

[77] Štrumbelj E, Kononenko I. Explaining prediction models and individual predictions with feature contributions. Knowledge and information systems. 2014;41(3):647–665.

[78] Lundberg SM, Lee SI. A unified approach to interpreting model predictions. Advances in neural information processing systems. 2017;30.

[79] Lundberg SM, Erion G, Chen H, DeGrave A, Prutkin JM, Nair B, et al. From local explanations to global understanding with explainable AI for trees. Nature machine intelligence. 2020;2(1):56–67.

[80] Pierce BG, Wiehe K, Hwang H, Kim BH, Vreven T, Weng Z. ZDOCK server: interactive docking prediction of protein–protein complexes and symmetric multimers. Bioinformatics. 2014;30(12):1771–1773.

[81] Sircar A, Chaudhury S, Kilambi KP, Berrondo M, Gray JJ. A generalized approach to sampling backbone conformations with RosettaDock for CAPRI rounds 13–19. Proteins: Structure, Function, and Bioinformatics. 2010;78(15):3115–3123.

[82] Forli S, Huey R, Pique ME, Sanner MF, Goodsell DS, Olson AJ. Computational protein–ligand docking and virtual drug screening with the AutoDock suite. Nature protocols. 2016;11(5):905–919.

